# The long-term evolutionary potential of four yeast species and their hybrids in extreme temperature conditions

**DOI:** 10.1101/2024.09.27.615317

**Authors:** Javier Pinto, Rike Stelkens

## Abstract

Accelerating climate change and extreme temperatures urge us to better understand the potential of populations to tolerate and adapt to thermal challenges. Interspecific hybridization can facilitate adaptation to novel or extreme environments. However, predicting the long-term fitness effects of hybridization remains a major challenge in evolutionary and conservation biology. Experimental evolution with microbes provides a powerful tool for tracking adaption, across generations and in real time. We investigated the thermal adaptation dynamics of four species of budding yeast (*Saccharomyces*) and their interspecific F2 hybrids, for 140 generations under cold (5°C) and warm (31°C) conditions.We found significant variation in the evolutionary potential of species and hybrids, strongly determined by their natural temperature tolerance. The largest fitness improvements occurred in hybrids, with some populations nearly quadrupling in fitness in the cold environment, exceeding both parents in thermal adaptive potential. Reciprocal transplanting of evolved populations from the endpoint of evolution into opposite temperatures revealed that hybrids had greater resilience when challenged with sudden temperature shifts. Our results highlight that hybridization alters the fitness outcomes of long-term adaptation to extreme environments and may render populations more resilient to sudden environmental change, presenting both opportunities and challenges for conservation and sustainable agriculture.

## Introduction

Hybridization, the exchange of genetic material between divergent populations and species, can facilitate adaptation, increase diversity, and rescue populations from extinction because it instantly introduces new genetic variation throughout the genome (Abbott et al., 2013; Grant & Grant, 2019; Seehausen, 2004; R. B. Stelkens, Brockhurst, Hurst, & Greig, 2014).

Hybridization rates may rise as a consequence of accelerating climate change and globalization (Canestrelli et al., 2017; Chunco, 2014; King et al., 2015; McFarlane & Pemberton, 2019), posing both risks and opportunities for adaptive evolution and conservation (Brauer et al., 2023). This has intensified the need to understand and forecast the evolutionary responses of hybrid populations to temperature changes but so far, our knowledge about the long-term fitness consequences of hybridization is limited. This is partly because the focus has traditionally been on long-lived hybridizing species of conservation concern (e.g. birds and mammals) with small population sizes. Making hybrid crosses in these systems and using them in experiments spanning multiple generations is difficult, if not impossible. Thus, most experimental studies rely on a short-term exposure of organisms to temperature stress and responses are usually measured within a single generation only (e.g. Petrović et al., 2023; Schwartz et al., 2024; Strait et al., 2024), which does not reflect the long-term changes occurring in populations over evolutionary time scales. In an applied context, how do we assess whether supportive breeding is appropriate if we do not know how outbred or hybrid populations respond to temperature changes? To start answering this, we need more experimental studies that investigate how temperature affects the capacity of hybrid populations to adapt, across evolutionary time scales and ecologically relevant thermal conditions.

Here, we used experimental evolution with species of the budding yeast genus *Saccharomyces* to assess the long-term fitness evolution of interspecific hybrids in cold and hot temperatures. Yeast is well suited for this work for two main reasons. First, *Saccharomyces* yeast species vary widely in temperature preferences ranging from 15° to 35°C (Salvadó et al., 2011). Heat resistance and optimal growth temperatures are well known across the genus with some cryotolerant strains able to grow at 0°C, while some heat-resistant strains can grow up to 42°C (Belloch et al., 2008; Pérez-Torrado et al., 2015). Divergence times between species are estimated to be between 2 million and 20 million years (Borneman & Pretorius, 2014; Kellis et al., 2003; Shen et al., 2018), and thermal adaptation is suggested to be a deeply diverged trait (Abrams et al., 2021; Weiss et al., 2018). This provides interesting starting points for experimental evolution and a rich resource to investigate the long-term fitness consequences of hybridization at different temperatures. Second, yeast can reproduce asexually by clonal ‘budding’, allowing for experiments with large population sizes (up to 200M cells), but also sexually by mating, followed by meiosis and recombination, which allows for making controlled hybrid crosses between species despite their large evolutionary divergence (Bendixsen et al., 2022;R.Stelkens & Bendixsen, 2022). Hybridization also occurs naturally in the genus and has substantially contributed to its origin and evolutionary history (Marcet-Houben & Gabaldón, 2015; Peris et al., 2017; Steensels et al., 2021). Interspecific yeast hybrids have been isolated from diverse natural (D’Angiolo et al., 2020; Eberlein et al., 2019; Leducq et al., 2016) and industrial environments, e.g. from lager beer fermentation (Gallone et al., 2019; Gibson & Liti, 2015; Hutzler et al., 2023). Thus, hybridization is a relevant driver of diversification in *Saccharomyces*.

The four yeast species we used represent all major phylogenetic clades found in *Saccharomyces* (Bendixsen et al., 2021) and vary widely in temperature preferences. Two of the species (*S. kudriavzevii* and *S. uvarum*) are considered cold-tolerant (Libkind et al., 2011; Naseeb et al., 2017; Salvadó et al., 2011). The Baker’s yeast *S. cerevisiae* represents the other end of the temperature spectrum and is considered warm-tolerant. *S. paradoxus* is considered a temperature generalist that grows well in a wide range of temperatures. We made recombined (F2) interspecific hybrid crosses between these species and used experimental evolution across 140 asexual generations, to explore the fitness dynamics and evolutionary potential for thermal adaptation of the parent species and their hybrids at two temperatures (5±1°C and 31±1°C).

### Materials and methods

### Parental strains

We used the four yeast species *S. cerevisiae, S. paradoxus, S. kudriavzevii,* and *S. uvarum*, which are representative of all major clades in the *Saccharomyces* phylogeny (Bendixsen et al., 2021). *S. cerevisiae* and *S. paradoxus* are sister species, i.e. they are the most closely related species in the *Saccharomyces* phylogeny (ca. 2 million years of divergence). *S. kudriavzevii* has an intermediate genetic distance to *S. cerevisiae* (10-15 million years) and *S. uvarum* is the most distantly related (10-20 million years) (Peris et al., 2023). We used two strains of each species: *S. cerevisiae* (XLY011 and Y55, both MATα mating type), *S. paradoxus* (YPS744 and Q89.8), *S. kudriavzevii* (ZP591 and FM1110), and *S. uvarum* (CBS7001 and JRY9189). All non-*S. cerevisiae* parents were MATa mating type (**Figure 1A**).

**Figure 1.**
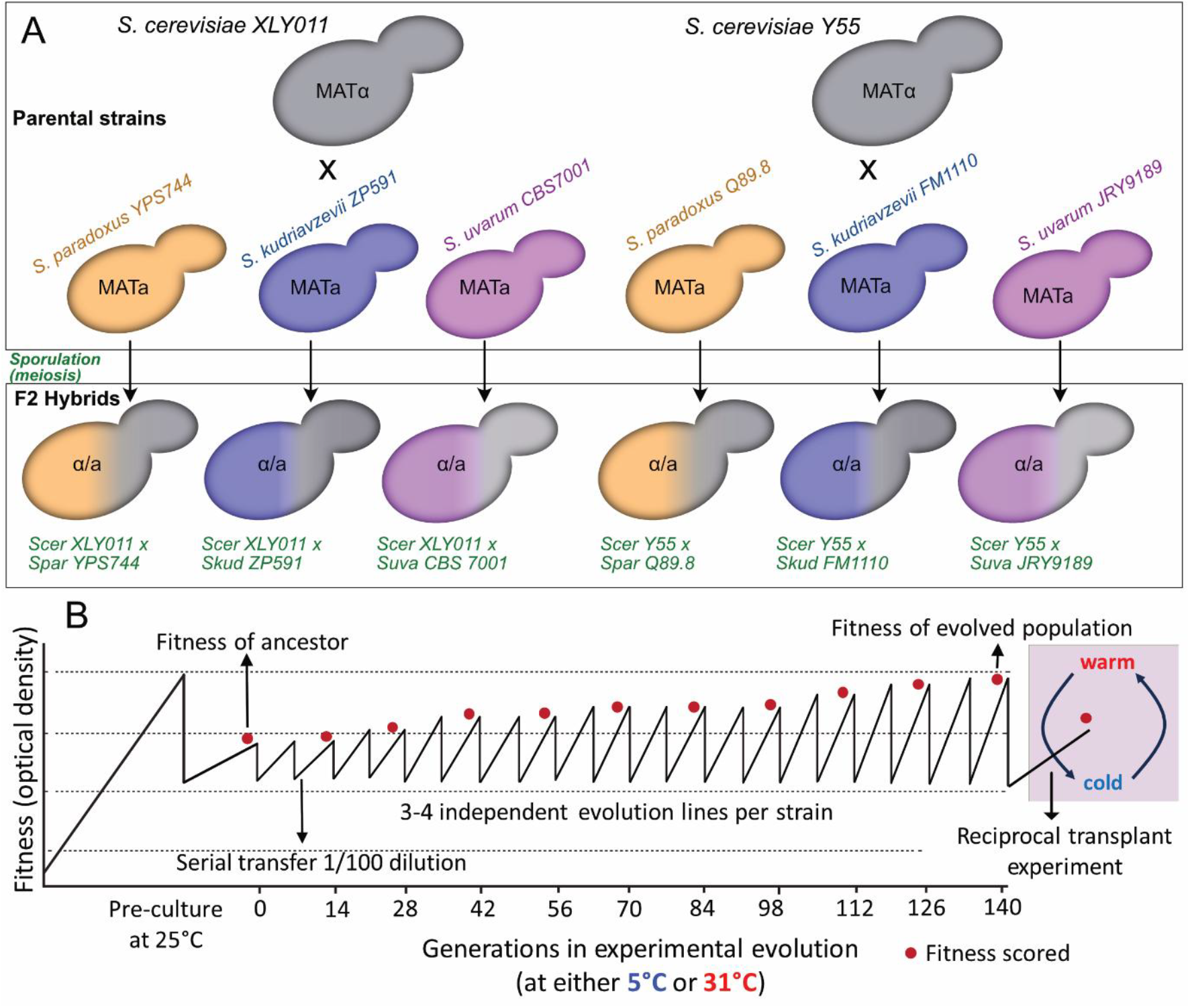
**A)** Crosses between two *S. cerevisiae* strains and strains of *S. paradoxus, S. kudriavzevii* and *S. uvarum,* resulting in six different F2 hybrids after sporulation (meiosis). **B)** Experimental evolution protocol with hypothetical fitness dynamics. Maximum biomass was measured as optical density (OD_600_) every 14 generations as a proxy of fitness (red dot), before serial transfer. Every population was evolved in warm (31°C) and cold (5°C) conditions, in three or four independent evolution lines (replicates), for ca. 140 mitotic generations. At the endpoint of evolution, a reciprocal transplant experiment was performed (purple square): The fitness of cold-evolved populations was measured in warm conditions and *vice versa*, after the sudden temperature shift.

Strains within species were isolated from different ecological niches (details in **Supplementary Table S1**). They differ genetically and phenotypically from each other (Liti et al., 2009; Peter et al., 2018; Warringer et al., 2011), but to different extents depending on the population structure and genetic diversity present within each species (ranging from only 0.4% sequence divergence within *S. cerevisiae* to ca 2% within *S. paradoxus*) (Peris et al., 2023). All strains used in this study were derived from diploid, non-domesticated isolates with the exception of *S. cerevisiae* XLY011, which is a homothallic diploid derived from BY4741 and BY4742 (both S288c) strain backgrounds commonly used in the laboratory. All strains were stable haploids, i.e. the mating-type switching endonuclease gene HO was inactivated by selective markers (either ho:KanMX, ho:HygMX, or ho:NatMX) to allow for targeted hybrid crosses.

### Construction of F1 and F2 hybrids

Haploid parents were grown overnight in liquid medium (YPD; 1% yeast extract, 2% peptone, 2% dextrose) in a shaking incubator at 30°C and then mixed at equal volumes.

Mating was induced by spinning down to concentrate cells and plating drops of 20μL on YPD plates (2.5% agar), kept at room temperature for 12h. We then streaked out for single colonies on a fresh YPD plate and incubated for 24 h at 30°C. Colonies were replica-plated to a sporulation plate (KAC; 2% potassium acetate, 2% agar) and incubated at room temperature for 5 days to induce sporulation. Diploids (F1 hybrids) were identified by inspecting individual colonies for tetrads (spore-containing asci) under the microscope and by testing for drug resistance to KanMX and NatMX to confirm the presence of the non-*cerevisiae* parent genome. A pure F1 diploid hybrid colony from each interspecies pair was then picked from the original YPD plate, propagated clonally in liquid YPD overnight, and frozen in 50% glycerol for later use. To obtain a genetically diverse, recombined F2 hybrid population, F1 hybrids were sporulated as follows. F1 hybrids were grown in liquid YPD overnight, cells were pelleted by centrifugation and transferred to a 500 mL Erlenmeyer flask containing 50 mL of liquid KAC, which was shaken at room temperature for seven days to induce meiosis. After microscopic verification of sporulation, the cultures were treated at 55°C for 30 min to kill any remaining unmated haploid (parental) and unsporulated diploid (F1 hybrid) cells (Rogers & Greig, 2009). Then, we centrifuged and washed cells, and resuspended them in liquid YPD to initiate germination and population growth. These large, recombinant F2 hybrid populations were frozen in 50% glycerol solution for later use.

To obtain six different hybrids, we used two strains of *S. cerevisiae* (XLY011 and Y55) and crossed each of them with one strain of the other three species, with increasing evolutionary divergence (*S. paradoxus, S. kudriavzevii*, and *S. uvarum*; **Figure 1A**). Thus, the two *S. cerevisiae* strains were used as reference strains with which all other strains were hybridized.

### Experimental evolution and fitness assays

We used experimental evolution to asexually propagate large diploid populations of all four species (eight strains in total, in three or four independent replicate evolution lines each) and six different interspecific F2 hybrids (in three independent replicate evolution lines each) at two temperatures; 5±1°C and 31±1°C (**Figure 1**). We chose 31°C for the hot evolution regime because this temperature still allowed the two cold-tolerant species (*S. kudriavzevii*, and *S. uvarum*) to grow. Note that replicate evolution lines of hybrid crosses are ‘biological replicates’ in the sense that each population contains genetically different individuals resulting from independent meiosis and recombination, while replicates of pure-species lines are technical, i.e. they were isogenic at the start of evolution.

To initiate experimental evolution, we inoculated 100μL of overnight cell culture (from frozen stock populations) into glass tubes containing 5 mL of liquid YPD. This corresponds to a starting population size of approximately 3 × 10^7^ cells. All populations evolved in the same incubator-shaker (one for each temperature) at the same time, i.e. they experienced the same slight temperature fluctuations if present. Continuous population growth was achieved by serially transferring 50μL of cell culture into new tubes with 5mL fresh YPD to provide nutrients after 24 hours at 31±1°C, and after 96 hours at 5±1°C. This corresponds to approximately seven population doublings in each temperature regime, between serial transfers. Generation time was calculated using the formula G= t / (3.3 log b/B) (G = generation time, B = number of cells at the beginning of time interval, b = number of cells at the end of time interval). We used a total of 20 serial transfers to reach approximately 140 generations in each temperature regime. Populations were standardized to the same cell density (optical density: OD_600_ = 0.5) before inoculating the next tube. This transfer protocol leads to periodic reductions in population size but bottleneck effects leading to potential loss of beneficial mutations are negligible, i.e. selection is stronger than drift in these systems (Wein & Dagan, 2019). To track the evolution of population fitness over time, we quantified maximum biomass of cells before each serial transfer using a spectrophotometer (Biotek Epoch2). We measured the OD_600_ of each population in three technical replicates, after two serial transfers, every 48 hours for warm-evolved populations and every 192 hours for cold-evolved, i.e. approximately every 14 generations. Ten OD readings were taken in total from each evolving population, spanning a total of 140 generations.

After 140 generations of experimental evolution, we reciprocally transplanted the warm-evolved populations (at 31±1°C) into the cold environment (5±1°C), and the cold-evolved populations (at 5±1°C) into the warm environment (31±1°C). We measured fitness following the same protocol and replication for OD readings as described above.

### Statistical analysis

Relative fitness (as a measure of the strains’ thermal evolutionary potential) was calculated by dividing the fitness of evolved populations at any number of generations by the initial fitness of their respective ancestor (i.e. before experimental evolution at generation 0). We used paired t-tests with Bonferroni correction to test for differences between ancestors and evolved populations and unpaired t-test with Welch correction to analyse differences between cold-and warm-tolerant species. The fitness of reciprocally transplanted populations was assessed in the same way, by comparing to the fitness of the respective ancestral strain.

We used a linear mixed effect model to test for potential sources of variation in relative fitness between parental species after experimental evolution. We used temperature (warm or cold), species (*S. cerevisiae, S. paradoxus, S. kudriavzevii, S. uvarum*), and the temperature-by-species interaction as fixed effects, and ‘strain’ and ‘evolution line’ as random effects. Similarly, to test for sources of variation in absolute fitness at the end of experimental evolution, we used a model with the initial absolute fitness of the strain (before experimental evolution), temperature (warm or cold), strain type (warm-tolerant or cold-tolerant), and the temperature-by-strain type interaction as fixed effects, and ‘strain’ and ‘evolution line’ as random effects. To test for sources of variation between parents and hybrids in relative fitness, we used temperature (warm or cold), cross type (parental or hybrid) and the temperature-by-cross type interaction as fixed effects, and ‘strain’ and ‘evolution line’ as random effects. All analyses were run in JMPv18.

## Results

We tracked the fitness dynamics and adaptation of four *Saccharomyces* species and their interspecific F2 hybrids using experimental evolution for 140 generations in a cold (5±1°C) and a warm environment (31±1°C; **Figure 2**). Across all strains, there was an inverse correlation between initial absolute fitness and relative fitness after experimental evolution (R^2^= 0.42; p < 0.01, **Supplementary Figure S1**), indicating that strains starting out with higher fitness showed smaller fitness gains over the course of the experiment. No correlation was found between initial and final absolute fitness (R^2^ = 0.005; p = 0.54).

**Figure 2.**
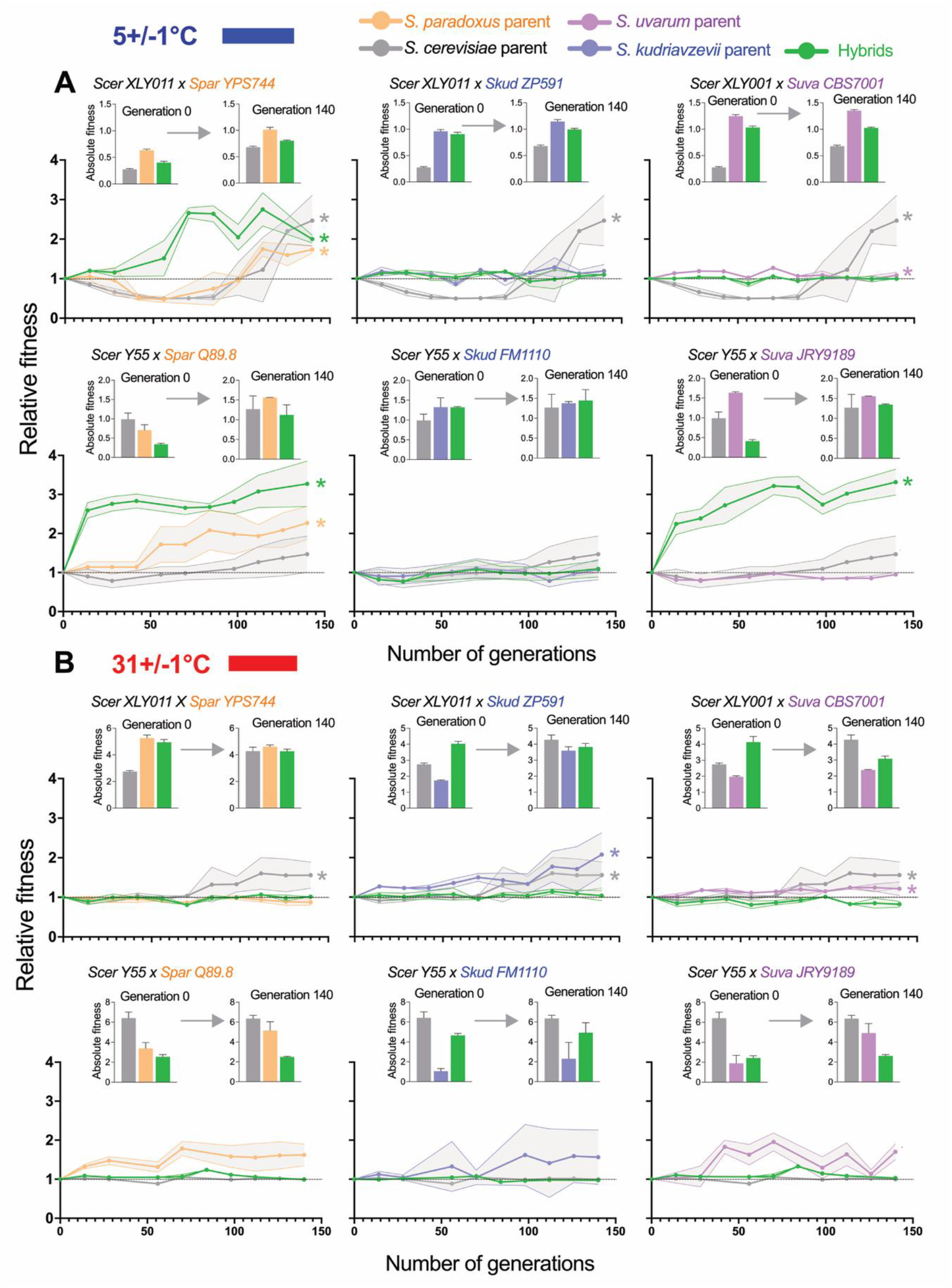
Relative fitness dynamics of eight strains of four *Saccharomyces* yeast species and six hybrid populations experimentally evolved for 140 generations in **A)** a cold (5±1°C) and **B)** a warm (31±1°C) environment. Lines show mean relative fitness across three or four independently evolved replicate populations per strain background. Grey-shaded areas show the standard deviations of replicates. Values above 1 represent fitness gains, values below 1 represent fitness losses, relative to the ancestral strain’s fitness. Asterisks indicate significant fitness gains from ancestral to evolved strains after 140 generations (results from paired t-test in **Supplementary Table S3**). Inserted bar plots show the absolute fitness of each strain before and after experimental evolution.

### Thermal evolutionary potential of parental *Saccharomyces* species

To compare the thermal evolutionary potential of the parental species, we tested for differences in relative fitness after experimental evolution (the fitness of evolved populations divided by the fitness of the ancestor). The model showed that temperature (warm or cold) and ‘species’ alone did not explain significant amounts of variance but the interaction terms between temperature and the species *S. cerevisiae, S. paradoxus*, and *S. kudriavzevii* were significant (t = 2.74, p = 0.011; t = −2.91, p < 0.01; t = 3.22, p < 0.01, respectively). This suggests that the evolutionary potential of the four species is affected differently by temperature. ‘Strain’ explained 26.23% of the variance (Wald p = 0.33) in relative fitness and ‘evolution line’ explained 2.61% (Wald p = 0.87; full model output in **Supplementary Table S2**).

Generally, strains of the warm-tolerant and generalist species (*S. cerevisiae* and *S. paradoxus*) showed larger fitness gains in the cold than in the warm environment. The cold-tolerant species (*S. kudriavzevii* and *S. uvarum*) on the other hand showed more fitness gains in the warm environment than in the cold. The largest improvements in relative fitness were found in *S. cerevisiae* at cold temperature (**Figure 2A**) and *S. kudriavzevii* at warm temperature (**Figure 2B**). *S. cerevisiae* XLY011 produced ~120% more biomass (averaged across evolution lines) than its ancestor at the end of evolution in the cold environment (t = 23.69, p = 0.002). *S. kudriavzevii ZP591* produced ~100% more biomass than its ancestor at the end of evolution in the warm environment (t = 13.32, p = 0.006; **Supplementary Table S3**). Together, these results suggests that the long-term evolutionary potential to thermally adapt depends on the natural temperature tolerance of the species, but also that considerable variation can occur between genetically different strains within species.

Next, we compared the relative fitness gains of the warm-tolerant and generalist species (*S. cerevisiae* and *S. paradoxus*) with that of the cold-tolerant species (*S. kudriavzevii* and *S. uvarum*) after experimental evolution. We found that warm-tolerant species generally showed higher thermal evolutionary potential at low temperatures (unpaired t-test with Welch correction: t_13,83_= 5.76; p < 0.001), whereas cold-tolerant species showed higher evolutionary potential at warm temperatures (t_16,45_ = 3.77, p= 0.007; **Figure 3**). Note, that as expected, cold-tolerant species (*S. kudriavzevii*, and *S. uvarum*) had higher absolute starting fitness than warm-tolerant species at cold temperature (**Supplementary Figure S2A, Supplementary Table S3**). At warm temperature, warm-tolerant species (*S. cerevisiae* and *S. paradoxus)* showed higher absolute starting fitness (**Supplementary Figure S2B**).

**Figure 3.**
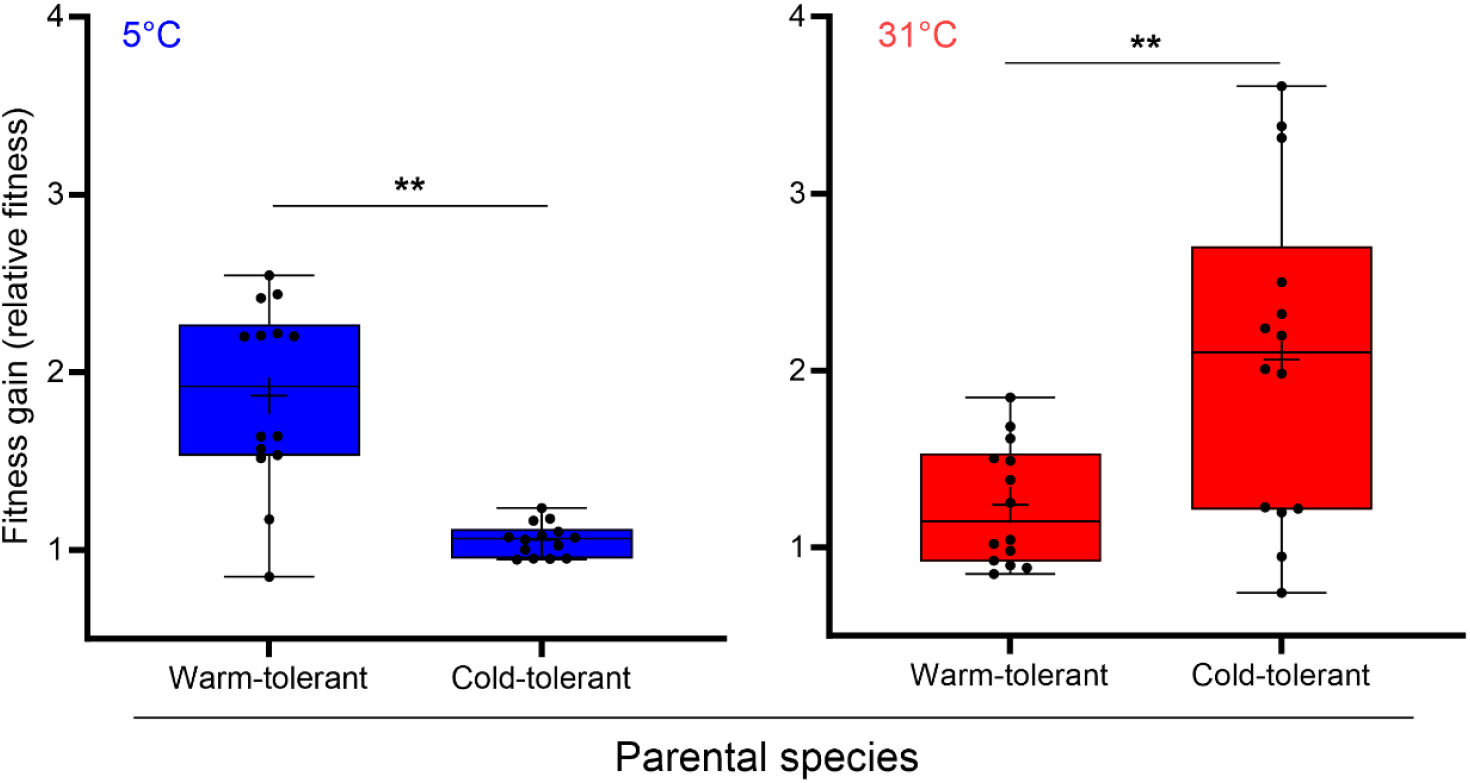
Comparing relative fitness gains after evolution in cold (5±1°C) and warm (31±1°C) environments between warm-tolerant and generalist (*S. cerevisiae* and *S. paradoxus*) and cold-tolerant (*S. kudriavzevii* and *S. uvarum*) species. Whiskers show the smallest and largest values, lines in the middle of the boxes represent the median and + shows the mean. Asterisks indicate significant differences (p < 0.01**).

Because of their noticeably different starting fitness, we also tested for sources of variation in absolute fitness gains between warm- and cold-tolerant species. The model testing for changes in absolute fitness (see inserted barplots in **Figure 2**) showed a significant effect of temperature (warm vs. cold) (t = −4.21, p < 0.01). The initial fitness of the strain (before experimental evolution) had a small effect, which was nearly significant (t = 2.04, p = 0.052). The thermotolerance type (warm- or cold-tolerant) and its interaction with temperature however, did not have significant effects on absolute fitness after experimental evolution, indicating that the temperature during experimental evolution did not differently affect the absolute fitness gains of strains with opposite thermal specializations. ‘Strain’ explained 20.5% of the total variance in absolute fitness change (Wald p = 0.3), while evolution line explained no variation (Wald p = 0.92; model output in **Supplementary Table S2).**

### Thermal evolutionary potential of interspecific hybrids

In the cold environment, three hybrid crosses significantly improved in relative fitness (**Figure 2A**). As opposed to their parents, the largest improvements in hybrid fitness were seen in the first 100 generations of experimental evolution, where both *S. cerevisiae* x *S. paradoxus* hybrid crosses, plus the *S. cerevisiae Y55* x *S. uvarum JRY9189* hybrid, exceeded both their parents in relative fitness. In the warm environment, hybrids fitness changed overall much less (**Figure 2B**) and the *S. cerevisiae XL011* x *S. uvarum CBS7001* hybrid cross lost fitness at warm temperature.

Hybrids often had low absolute starting fitness in the cold but improved over the course of evolution, with some populations doubling or tripling in absolute fitness (bar plots in **Figure 2A**). This was less pronounced in the warm environment, where the starting fitness of the hybrids was not as low as in the cold, but intermediate or similar to that of their parents (bar plots in **Figure 2B)**. Absolute fitness values and results of paired t-tests comparing evolved populations with their respective ancestors are shown in **Supplementary Figure S2** and **Supplementary Table S3.**

### Comparing the thermal evolutionary potential of parents and hybrids

When comparing the thermal evolutionary potential of parents and hybrids, no significant variation in relative fitness after experimental evolution was explained by cross type (parents *vs*. hybrids) alone. However, temperature (t = 2.41, p = 0.019) and also the interaction between temperature and cross type explained significant amounts of variation (t = 3.94, p < 0.01). ‘Strain’ explained 20.3% (Wald p = 0.047) of the variation and ‘evolution line’ explained no variation (Wald p < 0.01; **Supplementary Table S2)**. This suggests that the evolutionary potential of parents and hybrids is affected differently by the cold and warm temperature used during experimental evolution.

To visualize the fitness of parents and hybrids after experimental evolution in the warm and cold environment, we plotted the relative fitness gains at 31±1°C against the relative fitness gains at 5±1°C across all populations. We found that the largest improvements in fitness occurred in hybrids in the cold environment, with some populations nearly quadrupling in fitness (**Figure 4**). The majority of the parental populations (grey data points) also gained fitness at both temperatures, but not to the same extent as the hybrids. Confirming the patterns found above, hybrids showed the highest improvements at cold, particularly the two different *S. cerevisiae* and *S. paradoxus* hybrid crosses (orange data points, **Figure 4**) and the *S. cerevisiae Y55* x *S. uvarum JRY9189* cross (purple data points). Hybrid crosses between *S. cerevisiae* and *S. kudriavzevii* fell in the centre, showing no fitness gains in either the warm or cold environment (blue data points).

**Figure 4.**
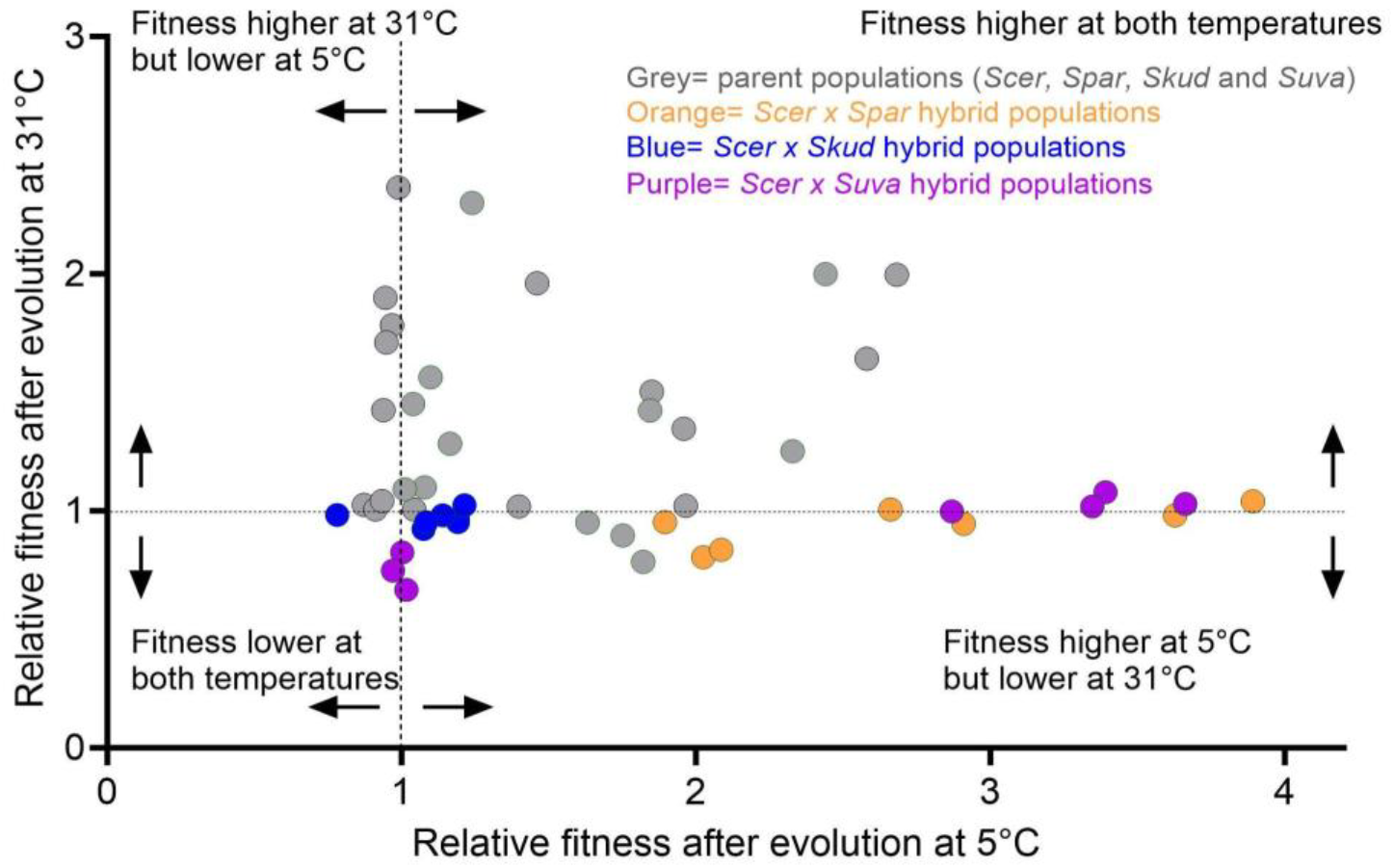
Relative fitness of parental strains and their six interspecific F2 hybrids after 140 generations of experimental evolution at 31±1°C, plotted against relative fitness after evolution at 5±1°C. Each data point shows the relative fitness of an independently evolved population (evolution line).

### Effects of sudden temperature shifts on the fitness of evolved parents and hybrids

After 140 generations of experimental evolution, we measured the fitness of all warm-evolved populations (at 31±1°C) in the cold environment (5±1°C), and the fitness of cold-evolved populations (at 5±1°C) in the warm environment (31±1°C), akin to a reciprocal transplant experiment. The transition from warm to cold caused significant fitness decreases in six of the eight evolved parent populations (**Supplementary Figure S3**). The two cold-tolerant species (*S. kudriavzevii* and *S. uvarum*) were most affected, with significantly lower fitness than the warm-tolerant species after the sudden temperature shift (ANOVA: F_2,11_ = 4.80; Tukey’s test: p = 0.025**; Figure 5A** left). One strain of the thermo-generalist *S. paradoxus* was unaffected while the other strain (YPS744) showed a significant fitness increase. The transition from warm to cold also significantly decreased the fitness of most hybrid crosses (**Supplementary Figure S3**). Overall, hybrid fitness after the temperature shift from warm to cold was intermediate between that of the parents, with no significant differences to either cold- or warm-tolerant parents (Tukey’s test vs. cold-tolerant: p = 0.23; vs. warm-tolerant p = 0.26).

**Figure 5.**
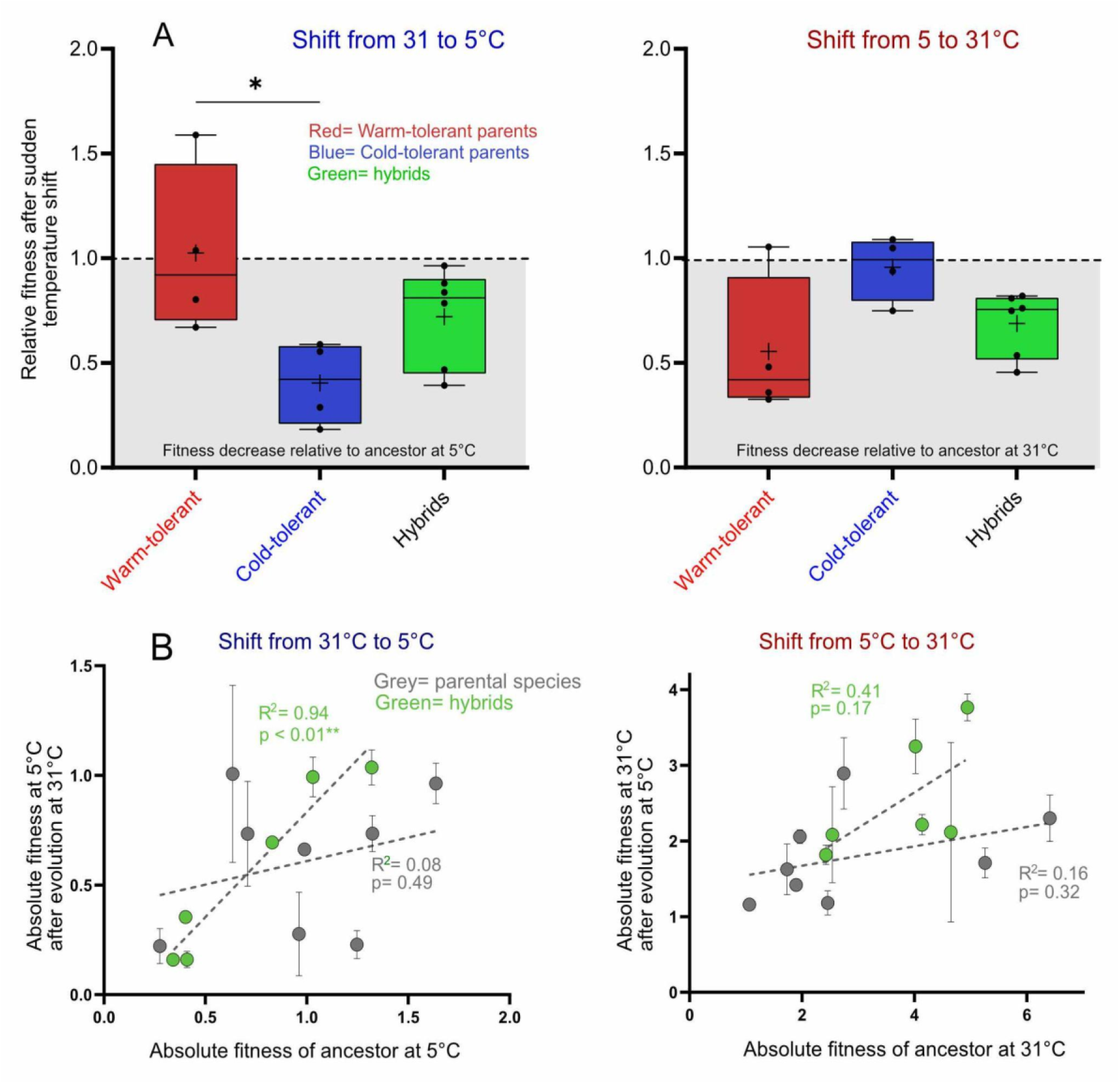
**A)** Left: The relative fitness of warm-tolerant species, cold-tolerant species, and their hybrids, after evolution in the warm environment (31±1°C), when suddenly exposed to the cold (5±1°C). Right: The relative fitness of these three groups after evolution in the cold environment (5±1°C), when suddenly exposed to the warm (31±1°C). Values above 1 represent fitness gains, values below 1 represent fitness losses, relative to the ancestral strain’s fitness. Whiskers show the smallest and largest values, lines inside boxes show the median and + shows the mean. Asterisk indicates significant difference in ANOVA. **B**) Linear regression of the absolute fitness of the evolved populations after sudden temperature shifts against the fitness of their ancestors. Asterisks indicate significant correlation coefficients.

The shift from cold to warm only caused significant fitness decreases in three of the eight evolved parent populations, of which only the two warm-tolerant species were affected (*S. cerevisiae* and *S. paradoxus*; **Supplementary Figure S3**). The fitness of the two cold-tolerant species (*S. uvarum* and *S. kudriavzevii*) remained mostly unaffected by the sudden temperature transition to warm. A comparison between the fitness of warm-tolerant and cold-tolerant species after the shift from cold to warm was non-significant (ANOVA: F_2,11_= 3.48; Tukey’s test: p = 0.06; **Figure 5A** right). However, the transition from cold to warm caused a significant fitness decrease in five of the six hybrid populations.

Again, hybrid fitness was overall intermediate between that of the parents, with no significant differences to either cold- or warm-tolerant parents (Tukey’s test vs. cold-tolerant: p= 0.19; vs. warm-tolerant p= 0.62).

We then tested if the original thermal performance of the strains before experimental evolution predicts how much their fitness was impacted by sudden temperature shifts, using regression analysis. We found that the absolute fitness of ancestral hybrid strains before evolution positively predicted their fitness after the temperature shift when transplanting populations from warm to cold (linear regression; R^2^= 0.94, p <0.01 **Figure 5B** left). Although we also found a positive trend when transplanting hybrids from cold to warm conditions, it was less strong and not significant (R^2^= 0.41, p= 0.17; **Figure 5B** right). For the parent species, there was no correlation between ancestral fitness and the temperature-shifted populations in either direction (R^2^ = 0.08, p = 0.49 in cold; R^2^ = 0.16, p = 0.32 in warm). This suggests that hybrids may have greater resilience when faced with sudden temperature shifts, especially when conditions change from warm to cold, and remain similar in fitness to their hybrid ancestor even after experimental evolution.

## Discussion

The effect of outbreeding and hybridization on thermal adaptation is of interest to evolutionary biologists, conservation managers, and agriculturalists alike. Genetic exchange in the form of gene flow, introgression and hybridization occurs in the majority of eukaryotic taxa and may become intensified through climate change and globalization (Chunco, 2014; McFarlane & Pemberton, 2019). Outbreeding across large genetic distances is usually a conservation concern because it can lead to offspring with poor fitness (Stelkens et al., 2015), maladaptation (Coyne & Orr, 2004), or produce more potent pathogens (King et al., 2015). But genetic exchange with divergent populations or species can also be a ‘saving grace’, particularly for endangered, genetically eroded populations that face stressful environments and require fast rates of adaptation to avoid extinction (Abbott et al., 2013; Brauer et al., 2023). However, predicting the long-term fitness effects of hybridization remains a major challenge in evolutionary and conservation biology to date.

Here, we used experimental evolution to measure the thermal adaptive potential of four *Saccharomyces* yeast species and their interspecific F2 hybrids in a cold (5±1°C) and a warm environment (31±1°C). Note that populations of parent species were isogenic at the start of our experiment, while hybrid populations were genetically diverse and potentially contained large amounts of genetic variation. Thus, the genetic basis of adaptation to temperature likely differs between parent and hybrid populations, which we discuss below.

We found that the yeast species we used as cross parents differed in their long-term evolutionary potential for thermal adaptation, depending on the temperature they were exposed to. In general, warm-tolerant species showed larger fitness improvements at cold temperature (**Figure 2A, Figure 3**) and cold-tolerant species showed larger fitness gains at warm temperature (**Figure 2B, Figure 3**), with some variation between strains within species. This pattern is consistent with predictions of diminishing-returns epistasis, limiting the size of available mutations with beneficial fitness effects in fitter genotypic backgrounds, which can constrain the rate of adaptation (Ardell et al., 2024; Chou et al., 2011; Wiser et al., 2013; Wünsche et al., 2017). Thus, strains already well-adapted to the temperature conditions used for experimental evolution showed smaller fitness gains.

When comparing the adaptive potential of the parent species against that of their hybrids, we found that some hybrids exceeded their parents in adaptation rates. Three of the six hybrids crosses showed a significantly higher potential to thermally adapt than both their parents. The most extreme fitness gains were observed in interspecific hybrids in the cold environment, with some populations nearly quadrupling in fitness relative to the fitness of their ancestral population before experimental evolution (**Figure 4**). The highest fitness gains occurred in a *S. cerevisiae* x *S. uvarum* hybrid cross and in both *S. cerevisiae* x *S. paradoxus* crosses. Similar to the parents, the largest fitness gains were observed in hybrid populations with low absolute starting fitness in the cold environment (lower than both parents individually; bar plots in **Figure 2A**). Note however, that at the end of the evolution experiment, hybrid populations did not usually attain a higher absolute fitness than the parents but rather converged to a similar fitness, making them at least competitive with the parents at the same temperature. Whether or not the adaptation rates we observed here would be sufficiently fast to allow hybrids to survive more extreme temperatures in the wild, where multiple abiotic and biotic stressors occur simultaneously (Rêgo et al., 2024), remains an open question.

While most parental populations showed fitness gains at both temperatures, most hybrids were constrained to fitness gains in cold temperature only. We speculate that this may be due to species-specific mitochondrial inheritance biases. Interspecific yeast hybrids contain two homologous genomes, but mitochondrial DNA is only retained from one of the parent species (Bendixsen et al., 2021). Which mitotype is retained depends on the selective pressure of the environment and determines, often to a large extent, the temperature tolerance of yeast hybrids (Baker et al., 2019), shaping their phenotypic and genomic evolution (Visinoni & Delneri, 2022). In our experiment, the fact that at cold temperatures, hybrids derived from warm-tolerant parents (*S. cerevisiae* x *S. paradoxus*) had a lower absolute starting fitness than hybrids with at least one cold-tolerant parent (**Supplementary Figure S2**), suggests that this is due to the lack of mitochondria from a cold-tolerant species. Consistent with this, *S. cerevisiae* x *S. uvarum* hybrids have been shown to retain mitochondria of the cold-tolerant S. *uvarum* (Hewitt et al., 2020) when grown at cold temperatures and *S. pastorianus* - a natural *S. cerevisiae* x *S. eubayanus* hybrid used for lager beer brewing under cold conditions - typically retains the mitochondria from the cold-tolerant parent *S. eubayanus* (Baker et al., 2019).

In our experiment, the increase of relative fitness in hybrids in the cold is likely a result of selection on the standing genetic variation produced by meiosis and recombination in hybrid populations, favouring the best recombinant genotype combinations for thermal adaptation, especially in crosses where no mitochondria from cold-tolerant species were present. In the parent populations on the other hand, which were isogenic at the beginning of experimental evolution, genetic adaptation was only possible through the (slow) acquisition of beneficial *de novo* mutations, which also explains their slower rise in fitness over generations (**Figure 2**). Thus, one may ask whether the comparison of parent vs. hybrid populations is evolutionarily relevant, since they have different genetic starting conditions. However, natural yeast populations typically contain very little genetic variation. Work on yeast isolates from the wild found high levels of clonality across space and time (Koufopanou et al., 2006) and often a complete absence of heterozygosity (Johnson et al., 2004; Sniegowski et al., 2002). Thus, we suggest that comparing the evolutionary rates of genetically diverse hybrid populations with that of genetically more uniform parent populations, does reflect, to an extent, the genetic conditions found in natural hybrid zones of yeast. For instance, in a narrow contact zone between the ranges of two nascent yeast species in North America, it has been suggested that hybridization has led to the formation of a new hybrid species, which has a potentially unique thermal niche that is different from both parental niches(Leducq et al., 2016).

Interestingly, in our experiment, most hybrids made from parent species with opposite thermal preferences showed no fitness gains in either warm or cold evolution conditions (except for *S. cerevisiae* Y55 × *S. uvarum* JRY9189 at 5°C; **Figure 4**). Especially the hybrid crosses between the naturally warm-tolerant *S. cerevisiae* and the cold-tolerant *S. kudriavzevii* did not improve in fitness, neither in the warm nor in the cold environment (**Figure 4**; blue dots). This is consistent with theory (Barton, 2001; Chevin et al., 2014; Rieseberg et al., 1999) and empiricism (Chhina et al., 2022; R. Stelkens & Seehausen, 2009) suggesting that F2 hybrids made from parents with a history of divergent selection for ecologically important traits (like thermotolerance) often display intermediate or ‘mismatched’ phenotypes, because there are fewer opportunities for complementation or epistasis of parental alleles with opposite signs at quantitative trait loci (which can otherwise result in transgressive traits). Adaptive responses of hybrid populations to environmental stress have been shown to be extremely variable (Brice et al., 2021; Gaworecki & Klaine, 2008; Lopandic, 2018; Oberhofer et al., 2014; Stelkens et al., 2014a; Stelkens et al., 2014b) suggesting that complex adaptations requiring changes in quantitative traits are often based on different genes in different species. This can lead to additive, dominant, and epistatic genetic interactions with different fitness effects in interspecific hybrids (Zhang et al., 2020). To generate more accurate predictions for hybrid fitness when exposed to thermal challenges, making crosses from parents with different genomic and ecological backgrounds can be highly informative, to which our study contributes.

Our reciprocal transplant experiment after adaptive evolution in opposite thermal conditions revealed interesting patterns. Sudden transitions from warm to cold had larger fitness costs than *vice versa*, especially for cold-tolerant species that had been forced to adapt to warm conditions (**Figure 5A** left). Shifts from cold to warm only negatively affected warm-tolerant species and thermo-generalists, which had been forced to adapt to cold conditions (**Figure 5A** right). These results suggest that beneficial mutations and molecular mechanisms leading to warm temperature adaptation may have deleterious fitness effects at cold temperatures (consistent with antagonistic pleiotropy). Alternatively, when evolving in warm environments, cold-tolerant species may accumulate loss-of-function mutations in genes normally needed for growth at cold temperatures, rendering them ill-adapted to their cold ‘home’ environment. The fact that the sudden shift from cold to warm left the two cold-tolerant species unaffected (*S. kudriavzevii* and *S. uvarum*), suggests that the genes and mechanisms underlying cold adaptation do not have detrimental effects on fitness at high temperatures. It is however important to note here that we exclusively used maximum biomass (yield) as a proxy for fitness. It is possible that including additional parameters across the full growth cycle, such as lag time, exponential growth rate, and stationary phase dynamics, would provide a more comprehensive understanding of temperature adaptation (Kaminski Strauss et al., 2019).

Interestingly, hybrids showed an overall greater resilience to sudden temperature changes than the parent species, especially from warm to cold conditions (**Figure 5B**). Even after 140 generations of evolution under opposite temperature conditions, hybrids still showed a similar performance as their ancestors: hybrids initially growing well at 5°C still grew well at 5°C, even after evolution at 31°C; and hybrids initially growing well at 31°C still had high fitness at 31°, even after evolution at 5°C. Recombinant hybrid genetic backgrounds may allow for better (or faster) compensation of sudden environmental shifts than the non-recombined parental genomes, for instance through activation of stress-response proteins (Nisamedtinov et al., 2008) or the altered expression of parental genes (Runemark et al., 2024). Gene expression analysis of ortholog genes in a domesticated hybrid yeast used for lager beer brewing (*S. pastorianus*) at cold temperatures showed that alleles inherited from the cold-tolerant *S. eubayanus* parent were significantly over-expressed compared to the warm-tolerant *S. cerevisiae* alleles (Timouma et al., 2021).

Alternatively, hybrid genomes may have more limited access to adaptive *de novo* mutations over the course of experimental evolution, due to lower rates of loss of heterozygosity (LOH), which can result from gene conversion or mitotic crossover in yeast (Smukowski Heil, 2023). LOH can allow beneficial mutations, even when recessive or only partially dominant, to sweep to fixation quickly in homozygous parental genomes, but may be masked by heterozygosity in the hybrids, as recently demonstrated in yeast hybrids challenged to evolve in an environment containing a UV mimetic drug (Bautista et al., 2023). This is consistent with our finding that hybrids adapted more slowly (or to a lesser extent) than the parents to the warm environment (**Figure 2B**). Thus, upon temperature reversal and exposure to the cold, hybrids may be better equipped to still grow well in the cold ‘ancestral’ environment because their ‘cold-adapted alleles’ were preserved.

## Conclusions

Genomic studies on vulnerable populations lacking the genetic variation to adapt and persist to climate change, typically ignore hybridization as a source of novel adaptive variation. But recent work on natural hybrids between ‘climate generalist’ species and species with more narrow climatic ranges (‘climate specialists’) suggests that hybrids have reduced vulnerability to environmental change, due to the fitness benefits of adaptive introgression (Brauer et al., 2023). Our findings confirm that hybridization alters the fitness outcomes of long-term adaptation to extreme environments and can contribute to population resilience when challenged by sudden temperature changes. Experimental evolution can provide relevant information for the conservation of genetically eroded, climate-sensitive populations under consideration for supportive breeding, but also for the breeding of climate-resilient hybrid crops with improved abiotic stress resistance (Cooper & Messina, 2023; ter Steeg et al., 2022). However, more studies on hybrids with different genetic and ecological parental backgrounds under different climatic conditions are needed if we want to understand what determines the long-term adaptive outcomes of hybridization.

## Author contributions

J.P. conducted the experimental and analytical work and drafted the figures. J.P. and R.S. jointly wrote the manuscript.

## Acknowledgements

We thank Chloé Haberkorn and Jennifer Molinet for their constructive comments on an earlier version of the manuscript.

