## Supplementary Figure S2 for "The long-term evolutionary potential of four yeast species and their hybrids in extreme temperature conditions"

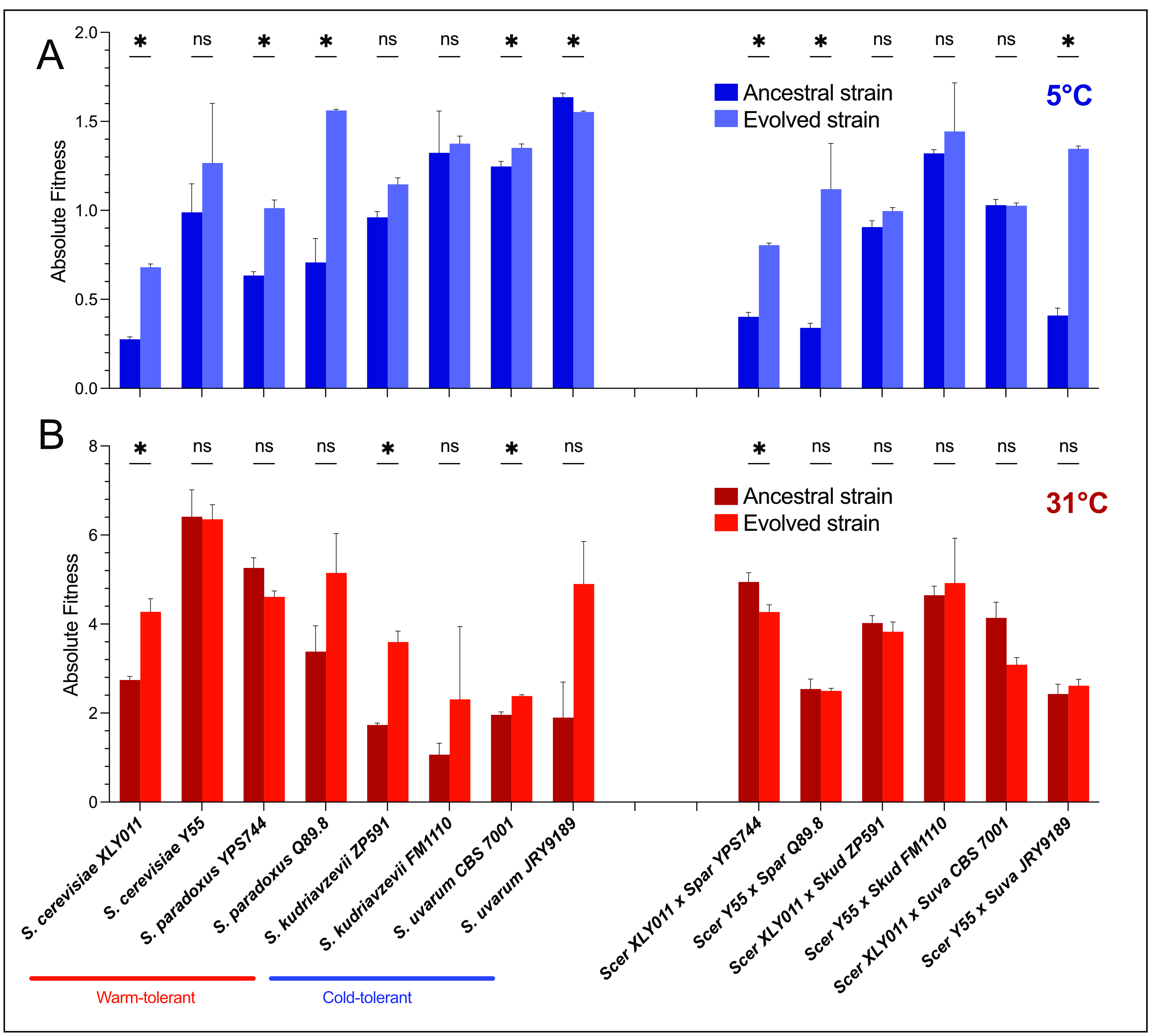

***Supplementary Figure S2.***  *Absolute fitness of ancestral and evolved strains and their hybrids, evolved at 5±1°C (panel A, blue) and at 31±1°C (panel B, red). Dark colour shows ancestral strain fitness (generation 0), light colour shows evolved strain fitness (at generation 200). Strains with names in red are considered thermo-tolerant or thermo-generalist, strain with names in blue are considered cold-tolerant. Asterisks show significant pairwise differences between ancestors and evolved populations after correction for multiple testing. Note the difference in y-axis scale in A) and B).*
