## Supplementary Figure S3 for "The long-term evolutionary potential of four yeast species and their hybrids in extreme temperature conditions"

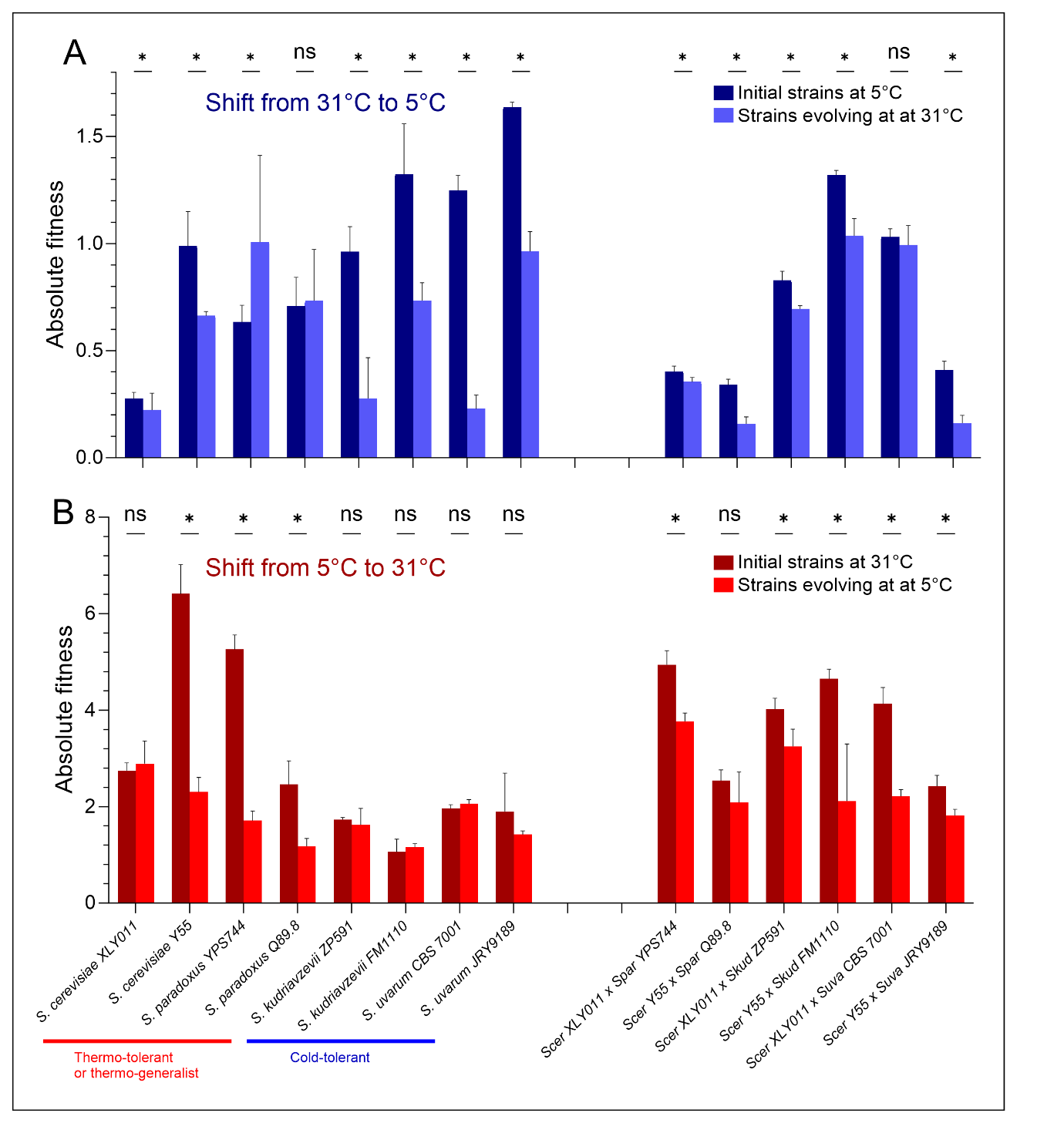

***Supplementary Figure S3.*** *Results of reciprocal transplant experiment of all parental strains (phylogenetic tree shown) and their interspecific hybrids.* ***A) Transition from warm to cold****: Blue bars show the initial absolute fitness of all populations (before evolution) at 5±1°C. Light blue bars show the fitness of populations evolved for 200 generations at 31±1°C when tested at 5±1°C.* ***B) Transition from cold to warm:*** *Red bars show initial fitness at 31±1°C, light red bars show the fitness of populations evolved at 5±1°C, when tested at 31±1°C. Bars represent averages from three or four independently evolved populations per strain. Error bars indicate standard deviation (SD). Asterisks show significant pairwise differences between ancestors and evolved populations after pairwise Tukey posthoc tests and Šídák's correction for multiple testing. Note the difference in y-axis scale in A) and B).*
