## Supplementary Figure S3 for "The long-term evolutionary potential of four yeast species and their hybrids in extreme temperature conditions"

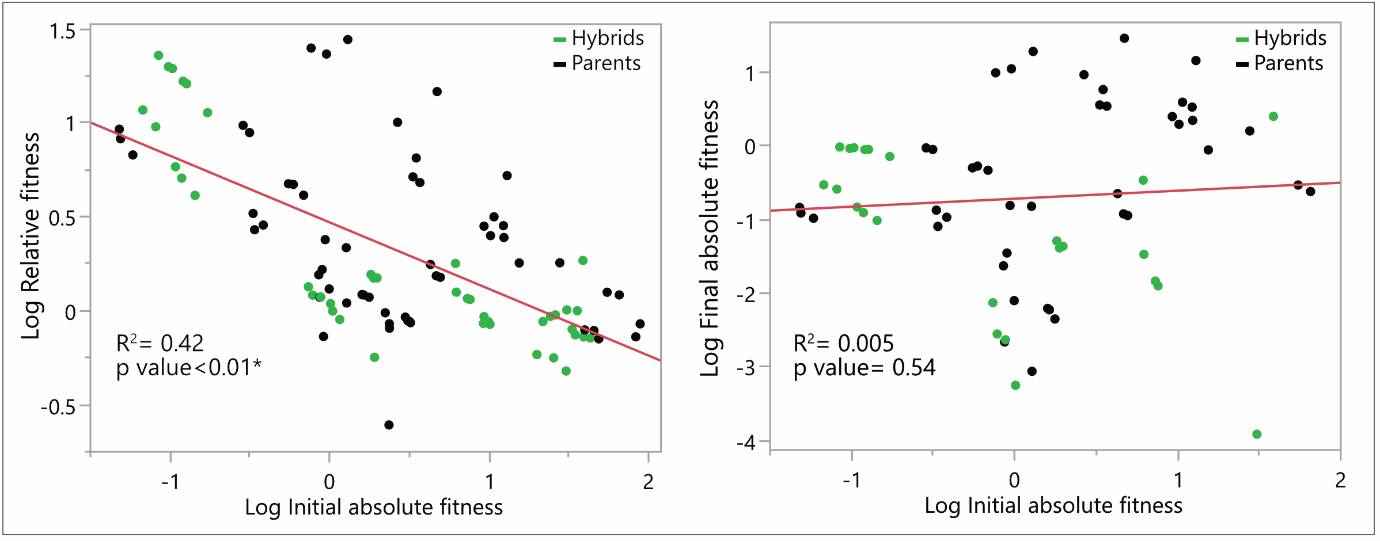

***Supplementary Figure S1.***  *Regression between initial absolute fitness and relative fitness after experimental evolution (left) (R^2^= 0.42; p< 0.01*) and regression between initial absolute fitness and final absolute fitness (right) (R^2^= 0.005; p= 0.54). Black dots show parental strains; green dots show hybrid strains.*
